# Thermogenic genes are blunted whereas brown adipose tissue identity is preserved in human obesity

**DOI:** 10.1101/2020.05.07.082057

**Authors:** Naja Z. Jespersen, Maja W. Andersen, Verena H. Jensen, Thit W Stærkær, Mai C.K. Severinsen, Lone Peijs, Ricardo Soares, Isabel Forss, Eline S. Andersen, Christoffer H. Hahn, Preben Homøe, Susanne Mandrup, Bente K. Pedersen, Søren Nielsen, Camilla Scheele

## Abstract

Obesity associates with a reduction in cold-induced glucose tracer uptake in brown adipose tissue in humans, suggesting loss of thermogenic capacity. We therefore hypothesized that a whitening of BAT occurs in obesity and assessed the molecular characteristics of deep neck BAT in a cohort of 24 normal weight, 24 overweight and 22 obese individuals in comparison with subcutaneous abdominal white adipose tissue (WAT). We found that the major marker of BAT thermogenesis, *UCP1*, was associated with central but not general obesity. We performed transcriptomic analysis of BAT in a cohort of 27 individuals classified as normal weight, over-weight or obese, and additionally four subjects with type 2 diabetes (T2DM), dispersed among the 3 BMI groups. We identified 3204 differentially expressed genes between BAT and WAT in samples from normal weight individuals, including genes involved in thermogenesis, but also revealing differences in developmental and immune system related genes. In BAT from individuals with overweight or obesity, 202 genes were downregulated and 66 of these were involved in cellular respiratory pathways, likely reflecting previously observed reduction in thermogenic function with obesity. Importantly, most BAT selective genes were *not* affected, and isolated adipose progenitors differentiated into thermogenic adipocytes with equal frequency regardless of BMI group. In conclusion, our data suggest a retained BAT identity, with a selective reduction of thermogenic genes, in human obesity.

## Introduction

Within the last decade, activation or recruitment of human brown fat has gained considerable interest as a promising treatment strategy against obesity and associated conditions including the metabolic syndrome and type 2 diabetes^1–5^. Although heat production is the main function of BAT in human infants and small mammals, a secondary role in energy homeostasis and weight-balance is established in rodents^6^, and evidence supporting a similar role in humans is accumulating^1,3^. Since the establishment of functionally competent BAT in adult humans, several studies have examined the role of active BAT in energy homeostasis and metabolism in human adult populations^7–13^. These studies have estimated the amount of active BAT by measuring BAT glucose uptake following cold-induced sympathetic activation by using 18F-fluorodeoxyglucose (18F-FDG) positron-emission-tomography / computed tomography (FDG PET/CT)-scans^14^. One of the challenges with this approach is that it is not possible to distinguish non-activated BAT from metabolically inactive white fat (WAT)^15^. Thus, it is unclear whether the reduced BAT activity observed in obesity^16,17^, is related to a loss of, or whitening of BAT, or if the tissue rather enters a thermogenically dormant state which is no longer acutely responsive to sympathetic activation.

Interestingly, it has been proposed that adult human BAT is best described as beige/brite fat^18,19^, i.e. a cold-inducible type of thermogenic tissue mainly found in the inguinal depot of rodents acclimated to a cold environment^20,21^. The link to humans is based on initial observations of the heterogenous morphology of human BAT, composed of a mixture of unilocular and multilocular adipocytes^22,23^ and the expression of murine beige fat markers in adult human BAT samples^19,22^. However, recent studies have reported a similarly heterogenous morphology in “classical” BAT depots of mice when housed in a thermoneutral environment^24,25^. Furthermore, some studies have addressed the characteristics of dysfunctional rodent BAT in genetic and diet induced models of obesity, sometimes referred to as ‘whitening’ of BAT^26,27^. These observations suggest that BAT might be equally as dynamic as WAT by acquiring WAT-like properties in response to certain environmental stimuli such as reduced sympathetic activation.

We here investigated the molecular signature of the progressive, obesity-dependent decrease in BAT activity^16,17^ and assess the relation between the thermogenic BAT marker *UCP1* and measures of metabolic health. To further address whether BAT identity was lost in obesity, we analyzed the thermogenic capacity of adipose derived progenitors from subjects with and without obesity.

## Results

### Subjects

32 patients (33 for cell-samples) (26 women and 7 men) scheduled for operation due to benign thyroid disease were enrolled in the “in depth” study, which included perioperative collection of supraclavicular deep neck fat and abdominal subcutaneous fat and a pre-surgical metabolic characterization. The participants had a wide range in age (median; 55 years range; 23 - 74 years) and BMI (median; 27.4 kg/m^2^, range; 17.6 - 38.4 kg/m^2^) (see **Table 1** for subject characteristics of the entire cohort) but were incidentally evenly distributed between the BMI categories as defined by the World Health Organization (WHO); normal weight (BMI: 18.5-24.99; n=10), overweight (BMI: 25.0-29.99; n=11), and obese (BMI ≥ 30; n=11). As indicated above, one subject was underweight (BMI=17.6), but was included in the normal weight group. The pre-surgical characterization included blood sample analyses (**Table S1A**), a DXA-scan evaluating body-composition, and an oral glucose tolerance test (OGTT). Additional subjects (n=36) (37 for cell samples) agreed to partake in the perioperative collection of biopsies without the preoperative metabolic characterization. In these patients we obtained medical history, height, weight, waist circumference (WC) and hip circumference (HC) on the day of surgery. Thus, we obtained paired supraclavicular deep neck and abdominal subcutaneous fat biopsies from 68 (70 for cell samples) patients in total (**Table 1**).

**Table 1.** Subject characteristics.

|  | **Women (N=53)\*** | **Men (N = 15)\*** | **Total (N = 68)\*** |
| --- | --- | --- | --- |
| **Age (years)** | 51 (23 – 74) | 63 (34 – 70) | 54 (23 – 74) |
| **BMI (kg/m<sup>2</sup>)** | 26.6 (17.6 -39.2) | 27.1 (22.6 -42.9) | 26.6 (17.6-42.9) |
| **WC (Cm)** | 91 (51-123) | 102 (83 – 140) | 95 (61-140) |
| **WHR** | 0.84 (0.67 – 0.99) | 0.97 (0.8 – 1.1) | 0.87 (0.67 – 1.1) |
| **Primary Diagnoses** | Non-toxic goiter N = 46<br>Grave’s disease N = 5<br>Hashimoto’s disease N = 2 | Non-toxic goiter N = 12<br>Grave’s disease N = 3<br>Hashimoto’s diseaser N = 0 | Non-toxic goiter N = 48<br>Grave’s disease N = 8<br>Hashimoto’s disease N = 2 |
| **Selected chronic comorbidities** | DM II N = 1(+1) **<br>Hypertension N = 11<br>Hypercholersterolemia N = 3<br>CVD N = 0<br>AF N = 2<br>COPD N = 1 | DM II N = 2<br>Hypertension N = 1<br>Hypercholersterolemia N = 3<br>CVD N = 2<br>AF N = 2<br>COPD N = 1 | DM II N = 3(+1) **<br>Hypertension N = 12<br>Hypercholersterolemia N = 6<br>CVD N = 2<br>AF N = 4 |
|  |  |  | COPD N = 2 |
| **Selected medications** | $\beta_1$RB N = 6<br>Melatonin N = 2<br>Metformin N = 1<br>Insulin N = 0<br>Statins N = 3 | $\beta_1$RB N = 1<br>Melatonin N = 0<br>Metformin N = 2<br>Insulin N = 1<br>Statins N = 3 | $\beta_1$RB N = 7<br>Melatonin N = 2<br>Metformin N = 3<br>Insulin N = 1<br>Statins N = 6 |
| **MDTp (°C)** | 9.2 (-1.4 – 18.9) | 8.5 ( -1.4 – 18) | 9.2 (-1.4 – 18.9) |

Values are median and range. \*70 subjects; 54 women and 16 men participated altogether, but no tissue samples, only cell culture material was included for two of them, and they are not included in the current table. The primary diagnosis refers to the disorder for which the patient was operated. Non-toxic goiter is enlargement of the thyroid gland without associated hormone disturbances. BMI = Body Mass Index, WC = waist circumference, WHR = waist / hip ratio, CVD = cardiovascular disease (here minus AF), AF = atrial fibrillation, COPD = Chronic obstructive pulmonary disease, RB = receptor blocker, MDTp = mean daily temperature one week prior to surgery.\*\*One subject was not previously diagnosed with T2DM, but 2hr plasma glucose indicated diabetes, while HbA1c was normal.

### Decreased *UCP1* mRNA in human BAT is associated with increased abdominal obesity

*UCP1* mRNA expression was substantially higher in deep neck fat compared to subcutaneous adipose tissue (AT) (P < 0.0001) but displayed substantial variation between subjects. In accordance with previous observations^22^, some deep neck samples thus had very low expression of *UCP1*, resembling the undetectable or nearly undetectable *UCP1* expression observed in subcutaneous AT (**Figure 1A**). To validate *UCP1* mRNA as an indicator of for functionally competent BAT, we next investigated how it correlated with age, a consistently well-established determinant of BAT activity as estimated using PET/CT-scans in humans^16,28,29^. In accordance with these human in vivo studies, we found age to be inversely correlated with *UCP1* mRNA levels (P = 0.003, R^2^ = 0.13) (**Figure 1B**). Somewhat surprisingly, but possibly relating to the large variability in *UCP1* expression in non-obese subjects, *UCP1* expression was not consistently different between BMI groups and did not negatively correlate with BMI (**Figure 1C-D**). This was in contrast to what has previously been reported in studies using glucose tracer uptake during cooling as a marker of BAT activity^17,29,30^. A possible explanation for this discrepancy, is that the biopsies in our cohort were obtained at room temperature where variations in *UCP1* might not reflect thermogenic capacity to the same extent as when subjects are subjected to cold, as *UCP1* is highly induced in brown adipocytes upon norepinephrine stimulation^31,32^. However, most subjects having *UCP1* mRNA levels within the highest range were non-obese individuals. We therefore found it relevant to further explore the relations between *UCP1* mRNA data and the metabolic measurements that we had performed. Importantly BMI has limitations as a predictor of metabolic disease^33^, as it fails to distinguish between the disease-risk linked abdominal obesity and the protective gluteofemoral fat accumulation^34^. Therefore, waist circumference (WC) and waist/hip ratio (WHR) might be more relevant variables than BMI in the context of metabolically beneficial effects of BAT. We found a negative correlation of *UCP1* mRNA levels with both WC (P = 0.03, R^2^ =0.09) (**Figure 1E**) and WHR (P = 0.002 R^2^ =0.18) (**Figure 1F**). These findings are in agreement with previous PET/CT based studies demonstrating a negative relationship between BAT-activity and WC^35^, visceral fat mass^16,35–37^ and relative abdominal obesity measured as visceral/total fat mass^35^, although none of these studies have directly assessed relative upper versus lower body obesity such as WHR. We observed no relationship with hip circumference (data not shown). Due to the well-established relationship between age, sex, BMI and WHR, we next performed a multivariate regression analysis, including the above-mentioned variables as well as mean outdoor temperature one week prior to surgery, and applied manual backwards elimination of independent variables. In this model both age and WHR remained independently significant, whereas none of the other variables conferred additional explanatory strength (**Table S1B**). Of note, women in the study had significantly higher *UCP1* mRNA levels compared to the men (P = 0.008) (**Figure S1A**), but this is likely to be driven by age and WHR, which were both significantly lower in women compared to men (P = 0.021 and P < 0.0001 respectively, data not shown). Thus, our data suggests a link between increased abdominal obesity and decreased expression of the BAT activity marker *UCP1*. Based on the additional metabolic characterization which had been performed in some participants (N = 32 for tissue samples), we were able to further address the observed relationship between supraclavicular *UCP1* expression, body composition and metabolic traits. First, we sought to ensure that the associations observed in the full dataset were not markedly different in the smaller cohort. Despite the decreased statistical strength both age (P < 0.007, R^2^ = 0.20) and WHR (P < 0.004, R^2^ = 0.13) remained negatively correlated to *UCP1* (data not shown), whereas BMI was not correlated in this smaller sample set either, and neither was overall body fat percentage (BF %) (data not shown). Interestingly, and in accordance with the WHR association in the larger sample-set described above, there was a negative correlation between *UCP1* mRNA levels and android/gynoid fat ratio (AGFR) (P = 0.01, R^2^ = 0.20, **Figure 1G**) and a tendency for android fat percentage (P = 0.08, R^2^ = 0.10) (**Figure S1B**), emphasizing an association between deep neck brown adipose *UCP1* and overall fat distribution. In accordance with WHR and AGFR being predictors of increased risk of insulin resistance and diabetes^38^, deep neck brown fat UCP1 expression also correlated negatively with HBA1c (P = 0.04, R^2^ = 0.13, **Figure 1H**) and plasma glucose at the two hour time point (PGT120) of an oral glucose tolerance test (OGTT) (P = 0.02, R^2^ = 0.17, **Figure 1I**). There was a tendency towards a negative correlation with the homeostatic model assessment of insulin resistance (HOMAIR) (P = 0.08, R^2^ = 0.11) (**Figure S1C**) and fasting plasma glucose levels (FPG) (P = 0.07, R^2^ = 0.10) (**Figure S1D**), but the latter was driven by one subject with abnormally increased FPG. We found no correlation between *UCP1* mRNA levels and plasma lipids including plasma triglyceride or plasma markers of low-grade inflammation, such as high sensitive C-reactive protein (hsCRP) or absolute neutrophil count (ANC) (data not shown, see **Table S1A** for all plasma analyses for which an association with *UCP1* was investigated). Lastly, we performed a multivariate regression analysis again with manual backwards elimination of independent variables in this smaller, but more thoroughly characterized cohort, including the significantly correlated variables of the univariate analyses (AGFR was included instead of WHR), as well as BMI. The model with the highest degree of explanatory strength (adjusted R^2^ = 0.38) included age, AGFR and PGT120, although none of the variables remained independently significant (**Table S1C**). Overall, these results suggest that *UCP1* mRNA expression, as a marker of functionally competent BAT, is decreasing with age and is linked to glucose tolerance, possibly explained by a robust relationship with fat distribution.

**Figure 1.**
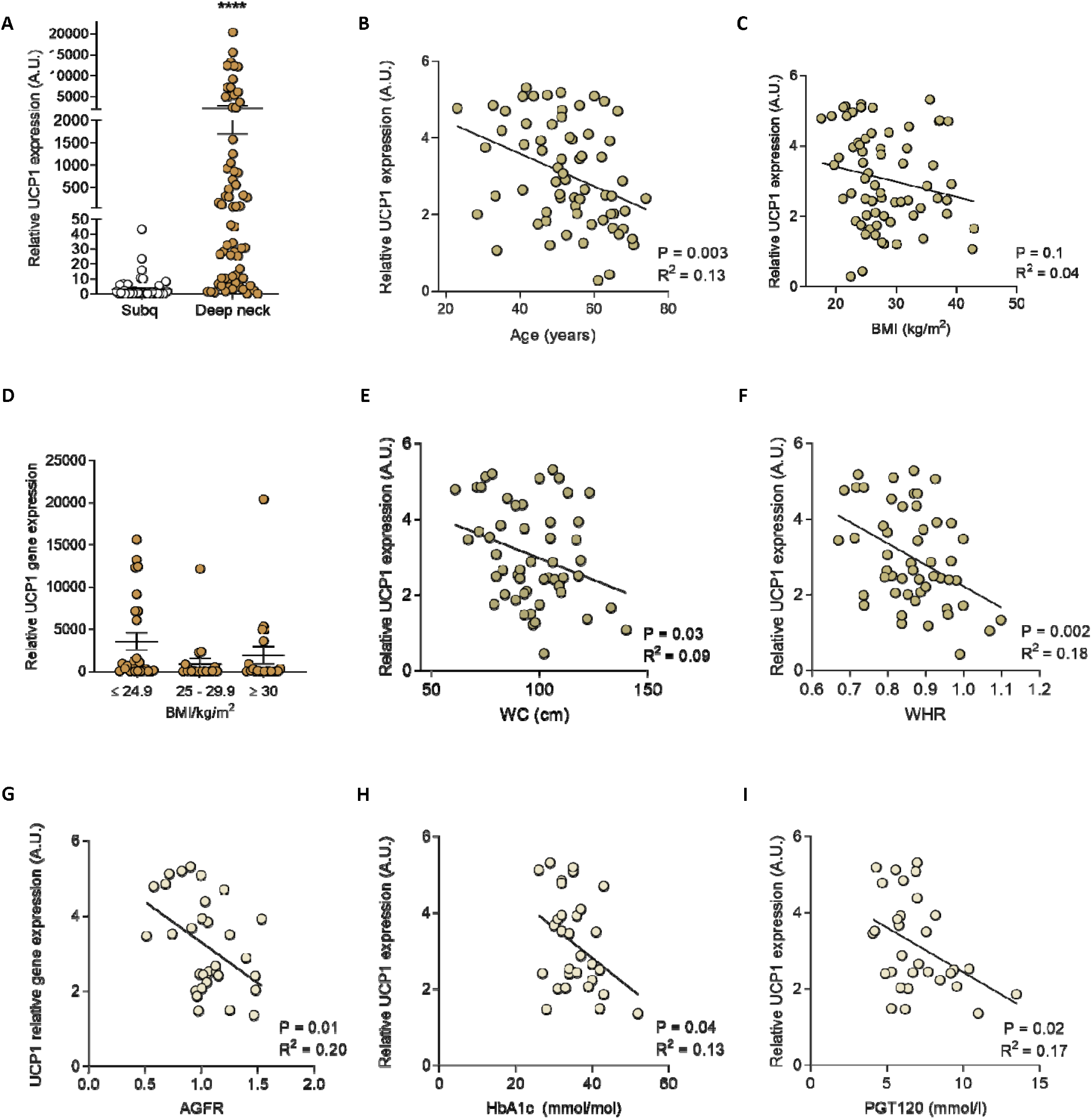
UCP1 expression in human BAT and association with metabolic characteristics. **A)** Relative UCP1 mRNA expression (mean + SEM) in subcutaneous abdominal WAT (N = 51) and supraclavicular BAT (N = 67). **B-C)** Correlations between log10 transformed supraclavicular UCP1 expression and **B)** age **C)** BMI. **D)** Relative UCP1 mRNA expression in supraclavicular BAT of NW (N = 25), overweight (N = 20) and obese (N = 22) patients (Mean and SEM). **E-H)** Correlations between log10 transformed supraclavicular UCP1 expression and **E)** waist circumference, **F)** waist / hip ratio (WHR), **G)** android / gynoid fat ratio (AGFR), **H)** glycated hemoglobin A1c (HbA1c) **I)** plasma glucose value at the 120 min time point of the OGTT (PG T120), AU = arbitrary units. Differences in gene expression between fat depots were assessed by Mann Whitney U test. The Kruskal–Wallis one-way ANOVA was used for comparisons between BMI groups. Univariate linear regression was performed for evaluation of correlations presented with unadjusted P-values. * = P < 0.05, **** = P < 0.0001.

### Comparative transcriptomics analysis of human brown and white adipose tissue

Given the negative correlation between BAT *UCP1* gene expression and central obesity, we aimed to investigate whether a “whitening” of BAT occurred in obesity. First, we determined the distinct gene signatures of human brown adipose tissue (BAT) and white adipose tissue (WAT), by performing RNA sequencing and comparative transcriptomic analyses. We analyzed a sub-set of BAT samples from the normal weight population of our cohort, obtained from deep neck fat in the supraclavicular area (n=7), and from subcutaneous fat in the abdominal area (n=4). Despite the previously observed heterogeneity of adult human BAT^22,31,39^ and the proposed similarity to the subcutaneously derived beige fat^19^, human BAT and WAT displayed distinct transcriptomes which were clearly separated as illustrated by a heatmap (**Figure 2A**). This separation between BAT and WAT was underscored with a principal component analysis (PCA) including all detected genes (**Figure 2B**). In confirmation of the BAT versus WAT integrity of the clustering, UCP1 was the number one gene contributing to PC1 (**Table S2A**). The plot further visualize and emphasize the heterogeneity of the BAT samples even within normal weight individuals, as has also been previously reported in BAT of adult humans^22^ (**Figure 2B**). We identified 3204 differentially expressed genes, with a false discovery rate (FDR) of 5%. Of these, 1239 genes were enriched in WAT, whereas 1965 genes were higher expressed in BAT. We assessed the most prominent WAT versus BAT selective genes by plotting the Log2 fold change and adjusted P-values (**Figure 2C**). As some genes displayed very low to undetectable expression levels in most BAT and WAT samples, and thus were considered unlikely to be intrinsic markers of one tissue type versus the other, we excluded genes that had an RPK (reads pr. kilobase) < 1 in samples of both BAT and WAT origin from this analyses. As a proof of concept, the thermogenic brown fat activity gene *UCP1*, displayed the strongest Log2fold upregulation in BAT versus WAT (**Figure 2C**). Also *LHX8*, which previously was described to discriminate between BAT and WAT in both human and mice studies^22,40^ were among the top BAT enriched genes (**Figure 2C,** and **Table S2B** for top 20 BAT enriched genes according to Log2foldchange and p-values). Only few of the top WAT enriched genes had previously been implicated in WAT or even adipose tissue function, possibly reflecting many similarities in mechanisms for lipid storage and lipolysis between BAT and WAT (**Figure 2C** and **Table S2C**). One gene higher expressed in WAT, with known adipose function, was *APOB*, encoding Apolioprotein B-100, which is involved in cholesterol metabolism^41^ while elevated plasma levels of this protein is a predictor of T2DM and is associated with WAT dysfunction^42^. Another WAT gene was *EGFL6*, which encodes Epidermal growth factor-like protein 6 a secreted protein previously reported to increase with obesity and to enhance proliferation of white fat precursor cells derived from human subcutaneous fat^43^. For comparison with previous studies, we plotted the expression of genes which were previously reported as markers for either WAT or BAT (**Figure 2D**). As expected, we observed a higher expression of *LEP*^44^ and *HOXC8*^22^ in WAT. In BAT, a variation between subjects was observed, but several markers in addition to *UCP1* and *LHX8* was enriched, including for example *EBF2*, *PRDM16*, *DIO2* and *CIDEA* (**Figure 2D**).

**Figure 2.**
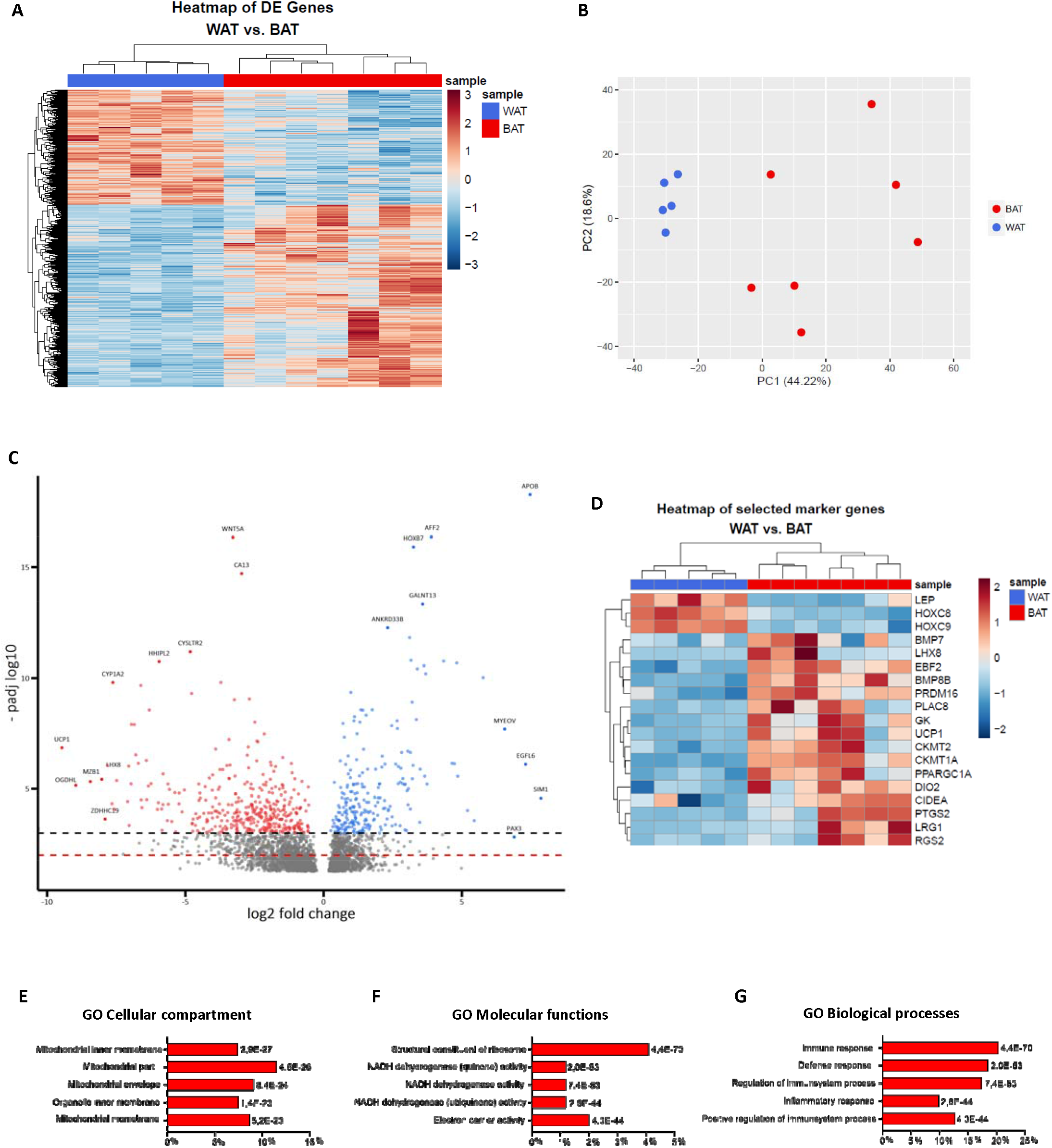
Comparative analysis of human BAT and WAT. **A)** Heatmap and **B)** PCA plot of all 3204 differentially expressed genes between BAT (n=7) and WAT (n=5) samples, FDR 5% **C)** Volcano plot depicting the top differentially expressed genes between BAT and WAT according to both P-value and log2fold change, top 10 in each category is specifically labelled. **D)** Heatmap of selected, established WAT (N = 5) and BAT (N= 7) marker genes. **E-G)** Top 5 gene ontology (GO) categories analyses of BAT enriched genes according to lowest p-values. **E)** Cellular Compartment **F)** Molecular functions **G)** Biological processes. X-axes depict percentage of enriched genes contributing to the respective GO-terms, P-values are presented for each category.

We performed a gene ontology (GO)-term analyses of the 1965 BAT versus WAT enriched genes. We found that the main BAT enriched cellular compartments were all related to mitochondrial structures (**Figure 2E**), and molecular functions to NADH-dehydrogenase and electron carrier activity (**Figure 2F**). This suggested a higher mitochondrial content and activity, a well-established property of BAT^6^. However, the GO terms for biological processes revealed a striking and somewhat unexpected BAT profile, since the top GO-terms were completely dominated by immune-related pathways and host defense mechanisms (**Figure 2G**), while processes related to cellular respiration and oxidative phosphorylation, although substantially enriched, were lower ranked. As indicated by the GO analysis, many of the BAT selective genes were related to various aspects of the immune system. One such gene among the top DE, which is also an established BAT-marker, is *IL6*, encoding the cytokine interleukin 6, which holds both inflammatory and anti-inflammatory properties^45,46^. *IL6* was one of the first recognized batokines^47^, i.e. secreted factors from BAT with endocrine, paracrine or autocrine functions, and has, at least in rodents, been demonstrated to be essential for some of the beneficial metabolic effects of increased BAT activity, particularly in regards to glucose homeostasis^48^. We further observed several immune related genes which have not previously been implicated in BAT biology (**Table S2B**). Some of these were related to B- or plasma cell function and antibody secretion; *MZB1*^49–51^, *IGLL5*, *FCRL5*^52^, *IGJ*^53,54^, and *CXCL13*^55^. Others are involved in leukocyte (neutrophil) function, adhesion, and migration; *SELE*^56–58^, *SELPLG*^59^, *CXCL1*^60,61^, chemokine activity and complement activation; *CXCL1*^62^, *CXCL13*^55^, and *PTX3*^63^, general inflammation and cytokine activity; *CXCL1*^61,64^, *PTX3, CYSLTR2*^65,66^ *WNT5A*^67^, T-cell activity and migration; *SELPLG*^59^, *CXCL13*^55^, and *WNT5A*^68^, direct antimicrobial activity direct antimicrobial activity and host defense; NTS69, LEPR70, and CXCL171, protection from viral entry into cells; PTX363, and macrophage activity; *CXCL1*^61^, *CYSLTR2*_66_, *WNT5A*^67^. Although immune cell infiltration in adipose tissue is often considered a hallmark of obesity related adipocyte dysfunction^72,73^, adipose tissue immune cells also exert explicit protective immune and anti-inflammatory functions under healthy conditions^74,75^, and is even directly involved in the defense against invading pathogens^76–78^. Of the top BAT enriched immune-related genes, some of them had previously been implicated in obesity related adipose tissue inflammation. This was the case for both of the chemokines CXCL1 and CXCL13, which exert both beneficial and detrimental effects depending on the context ^79,80,81,78^. Other top DE immune related genes have not previously been associated with a metabolic role in AT. An example of this is MZB1, which encodes the marginal zone B and B1 cell specific protein. MZB1 is localized in the endoplasmic reticulum (ER), where it functions as a cochaperone for another chaperone Grp94/Gp96 in circumstances of ER-stress. It has been demonstrated to be essential for the terminal differentiation of B-cells into antibody-secreting plasma cells, and specifically play an important role in the adaptive, T-cell-independent immune response^51^. It is a downstream effector of the PRDM1 transcriptional network in plasma cells, and involved in activation of the β_1_-integrin, which in turns is involved in mediating lymphocyte adhesion through binding to the vascular cell adhesion molecule (VCAM-1)^51^. In addition, it has been demonstrated to be of critical importance to IgA secretion and function in the gut mucosa, and thus mucosal immune defense and microbiotic homeostasis^50^. Of note, VCAM-1 was also among the BAT enriched genes, albeit not within the top-20 DE. All together the differential expression of these genes could indicate a specific immunoregulatory role of BAT, distinct from WAT. Many of the immune related genes, as well as other BAT enriched genes were encoding mitochondrial proteins, in line with a general higher mitochondrial content which is a well-established property of BAT^6^. Thus, although we found thermogenic genes and brown adipogenic markers to be higher expressed in BAT compared to WAT, the most prominent and novel finding was the diverse immunogenic profile in BAT compared to WAT.

### The deep neck brown fat gene signature in obesity

We next aimed to map the molecular signatures of human BAT in obesity and to assess whether BAT selective genes were downregulated in favor of WAT selective genes. We performed transcriptomic analysis of supraclavicular deep neck brown fat surgical biopsies from a cohort of 27 individuals including: normal-weight (n=7), overweight (n=10) and obese (n=10) subjects, as well as 4 patients with T2DM distributed over all three BMI groups (**Table S3**). The four patients with T2DM were excluded from the initial comparative analyses. We identified 896 differentially expressed (DE) genes in BAT when comparing the three BMI Groups (FDR 5%). To investigate our initial hypothesis of a whitening of BAT in obesity, as has been observed in rodents^26^, we performed an unsupervised clustering of the BAT and WAT enriched genes stratified for BMI and including the normal weight WAT samples (**Figure 3A**). Although all except one NW BAT sample clustered opposite to the WAT samples, the samples from overweight and obese subjects were intermixed between these two extremes with no clear BMI dependent pattern (**Figure 3A**). To further address our hypothesis, we performed a PCA plot including all three groups as well as the WAT samples from non-obese subjects, only including the genes that were differentially expressed between BAT and WAT (**Figure 3B**). However, this analysis suggested that the difference between BAT and WAT was maintained in obesity as none of the BMI groups clustered with WAT (**Figure 3B**). We found that 645 genes differed between samples from NW and OW individuals and 501 genes differed between NW and obese subjects, with 252 genes overlapping between the two conditions (**Figure 3C**). Among the overlapping genes 202 genes were downregulated in overweight and obesity and interestingly, 135 of these were higher expressed in BAT compared to WAT. Gene ontology (GO) analysis of the downregulated genes suggested involvement in oxidation-reduction, cellular respiration processes, and electron transport chain, all mechanisms important for thermogenic function (**Figure 3D**). Examples of mitochondrial genes that were downregulated with OW and obesity were *COX41A*, *COX8A* and *PDHA1* (**Figure 3E**). In contrast, these samples had upregulated expression of for example *LIMCHI*, *DMTN* and *NEIL2*, which are involved in cytoskeleton regulation and protein binding (**Figure 3F**).

**Figure 3.**
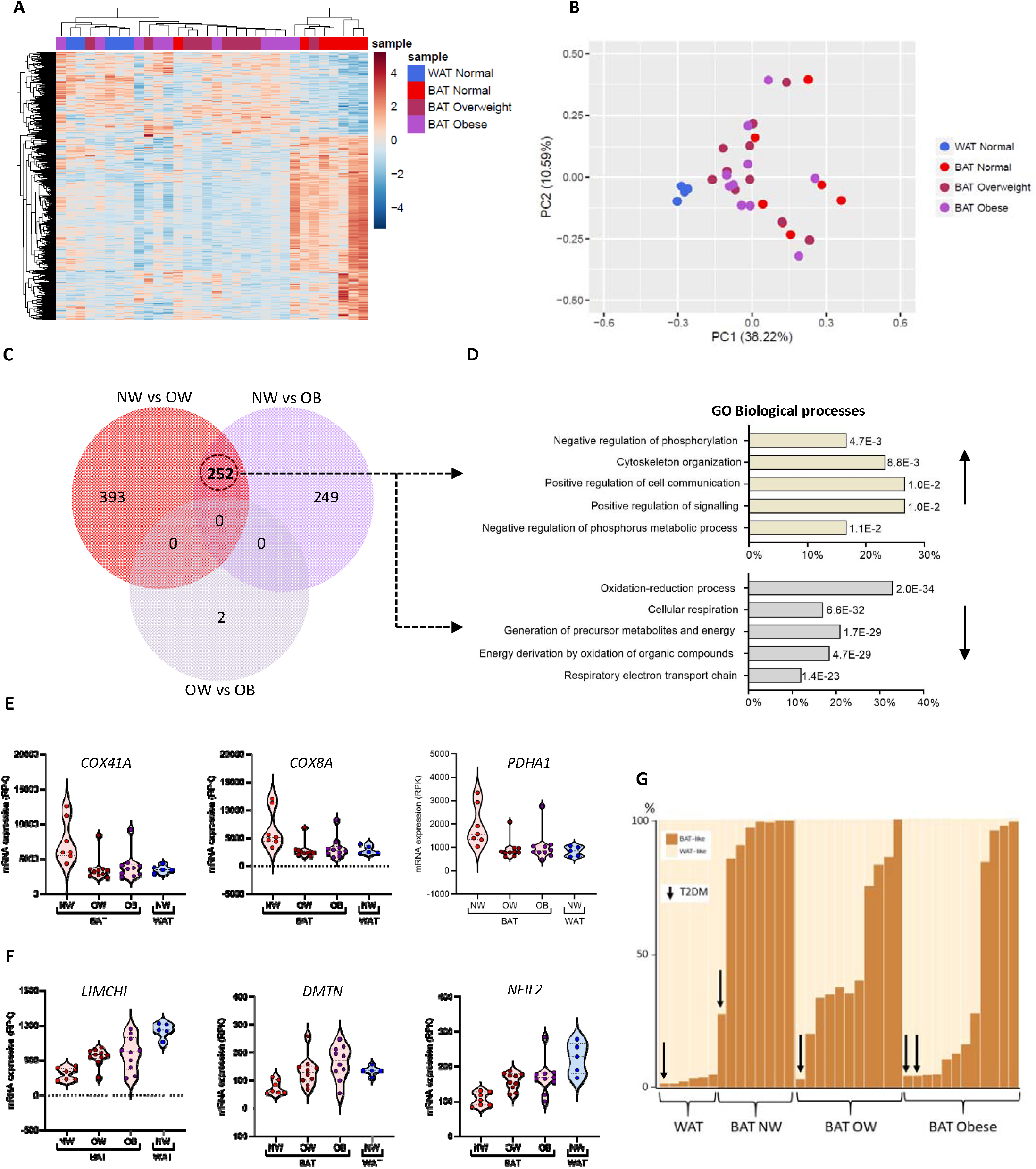
Transcriptomics of human BAT derived from normal weight, overweight and obese adult humans. **A)** Heatmap and **B)** PCA-plot of all DE expressed genes between WAT and BAT stratified according to BMI group. Normal weight BAT (n=10), Normal weight WAT (n = 5), overweight (n=11) and obese (n=11), using FDR 5%. **C)** Venn diagram depicting number of differentially expressed and overlapping genes in BAT between BMI groups. **D)** Gene ontology analysis, Top 5 biological process categories (BP), of the 135 BAT selective genes (upper-panel) which were down-regulated in BAT of overweight and obese compared to normal weight subjects, and the 30 WAT selective genes (lower panel), which were upregulated. X-axes depict percentage of enriched genes contributing to the respective GO-terms, P-values are presented for each category. **E)** mRNA expression of selected mitochondrial genes contributing to the top BP GO term categories of BAT selective genes downregulated in BAT of overweight (OW) and obese (Ob) compared to normal weight subjects (NW) **F)** and WAT selective genes involved in cytoskeletal regulation, up-regulated in BAT of overweight and obese compared to normal weight subjects. RPK = reads pr. kilobase. Data are presented as violin plots with Mean and SD. **G)** Percentage WAT and BAT like genes in individual biopsies from Normal weight WAT (n = 6), normal weight BAT (n=10), overweight BAT (n=11) and obese BAT (n=11), including 4 patients with type 2 diabetes (T2DM), as predicted by the computational tool PROFAT based on transcriptomic profiles.

These analyses suggested that despite an overall preserved BAT identity in obesity, the thermogenic function was reduced. To further validate these results, we used the validated computational tool ProFAT^82^, which can predict the thermogenic potential of a given sample set based on transcriptional profile^82^. (**Figure 3G**). As one separate bar is given for each sample, we here also included the four BAT samples and one WAT sample from patients with type 2 diabetes. The results are given and visualized as percentage WAT and BAT like genes in each sample (**Figure 3G**). As would be expected the subcutaneous WAT samples consisted almost entirely of WAT-like genes, whereas the normal weight BAT samples, despite the heterogeneity, nearly all expressed between 80-100% BAT-like transcriptomic profiles, a percentage which was substantially decreased with increasing BMI group (**Figure 3G**). Intriguingly, the samples originating from patients with T2DM, were predicted to contain the lowest amount of thermogenic capacity within each BMI group (**Figure 3G**), indicating a clear negative association between thermogenic BAT and glucose metabolism across BMI groups. Altogether, these results illustrate that while a thermogenic signature is reduced in BAT in subjects with overweight or obesity, BAT remains to be BAT with a transcriptome still distinct from that of WAT.

### Brown fat precursor cells are present in deep neck fat of adult humans with obesity

To further address the effects of obesity on deep neck brown adipose identity, we examined adipose derived progenitor cells from the described cohort. We have previously demonstrated that adipose progenitors derived from human supraclavicular brown adipose tissue differentiate into brown adipocytes, while adipose progenitors derived from a white adipose depot accumulate white adipocyte characteristics upon differentiation, despite application of the same culturing conditions^22,83^. Utilizing our previously optimized differentiation protocol with minor modifications as specified in the methods section, we thus differentiated adipose progenitors derived from either deep neck brown adipose tissue or from subcutaneous adipose tissue into mature lipid droplet containing adipocytes. Following exposure to either 1 μM of norepinephrine (NE) or vehicle treatment for 4 hours on the final day of differentiation cells were harvested for *UCP1* mRNA expression analysis. Deep neck-derived adipocytes had substantially higher *UCP1* mRNA levels compared to subcutaneous adipose-derived adipocytes even at baseline (P < 0.0001) and only deep neck adipocytes increased *UCP1* expression significantly following NE stimulation, (P < 0.0001) (**Figure 4A**). In line with what we observed in the tissue, cellular *UCP1* mRNA levels were highly variable between cultures from the deep neck depot, whereas all subcutaneous adipocytes expressed low basal *UCP1* levels and only very few displayed a modest NE-induction (**Figure 4A**). We observed no difference in the terminal adipocyte marker *FABP4* expression between deep neck and subcutaneous adipocytes (**Figure S4A**) or in visual estimation of lipid droplet accumulation (data not shown). This indicates equal lipid accumulation potential, and at the same time underscores that the higher *UCP1* mRNA expression in deep neck BAT compared to subcutaneous WAT cell cultures does not reflect a differentiation bias.

**Figure 4.**
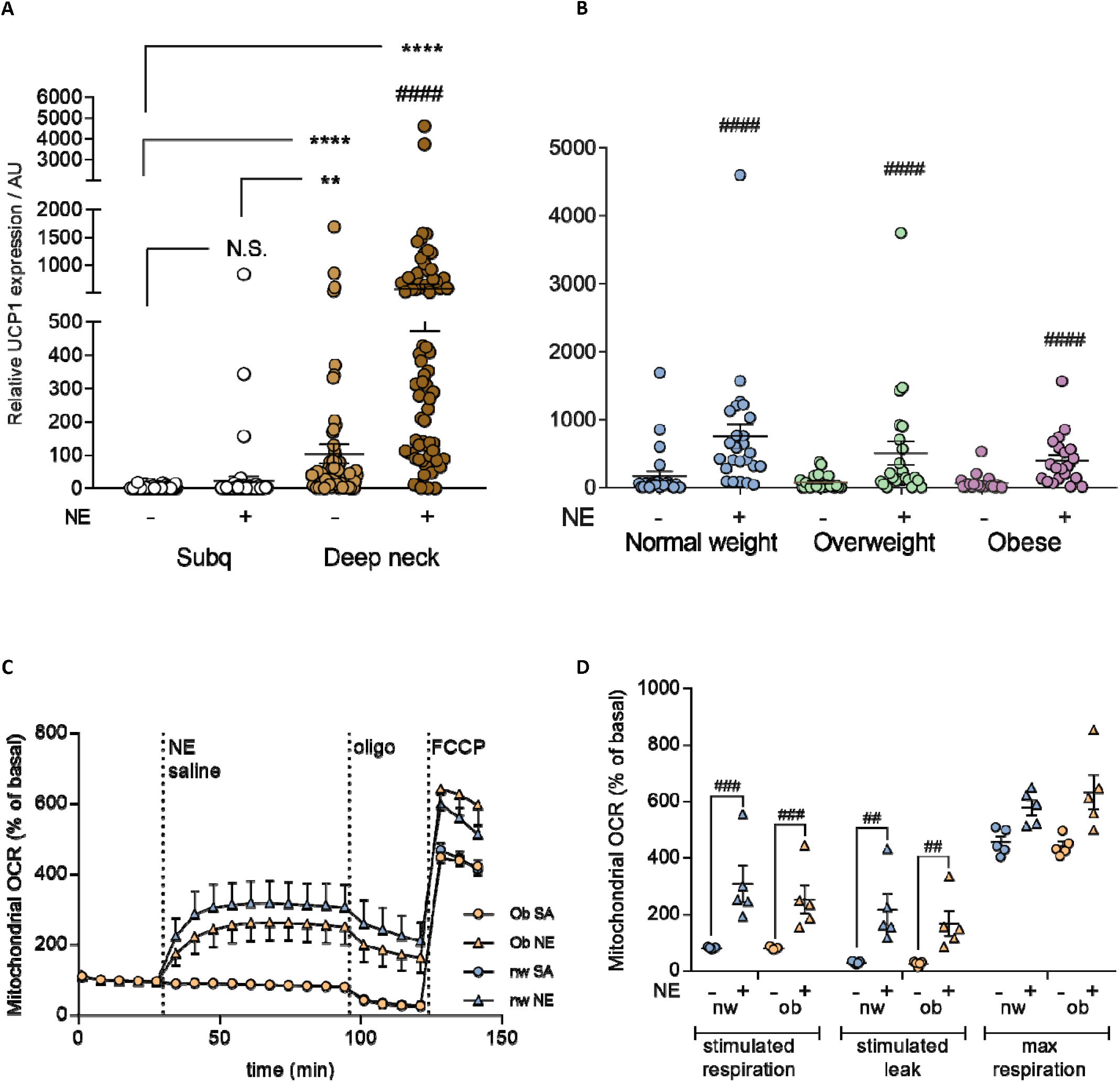
UCP1 expression and inducibility in primary human white and brown adipocytes. Relative UCP1 mRNA expression in paired primary subcutaneous adipocytes (WAT) and supraclavicular adipocytes (BAT). Data are presented as individual data points with line and whiskers representing mean and SEM. B) Relative UCP1 mRNA expression in paired primary supraclavicular adipocytes (BAT) with and without norepinephrine (NE) and divided into BMI groups. Data are mean and SEM, * = differences between depots, # = differences within depot +/− NE. # = P < 0.05, #### = P < 0.0001, **** = P < 0.0001. Differences in UCP1 expression between and within groups were assessed using repeated measures mixed models ANOVA with Bonferroni correction. A P-value below 0.05 was considered statistically significant.

We next wished to address a potential effect of obesity on cellular *UCP1* expression. Interestingly, deep neck cells displayed similar *UCP1* induction levels with NE stimulation across BMI groups (**Figure 4B**), i.e. *UCP1* was significantly increased with NE in both normal weight, overweight and obese participants (P < 0.0001), with no difference in basal or stimulated *UCP1* levels between groups (**Figure 4B**). In order to assess potential functional declines in brown adipocytes derived from obese subjects, we selected the brown fat precursor cells from the five donors, all women, with the highest BMI and matched them with cells from normal weight subjects, considering age, sex and differentiation capacity (i.e. estimated % lipid droplets) (**Table S4A**). Cells were differentiated in parallel, using the same differentiation protocol and norepinephrine-stimulated mitochondrial leak was recorded using the XF96 Seahorse Bioscience platform. In accordance with the results from the mRNA expression analysis, we detected no difference in the respiratory response between the brown adipocytes derived from obese subjects compared to brown adipocytes from the normal weight group (**Figure 4C, D**). In both groups, NE stimulated mitochondrial leak respiration, supporting thermogenic function (**Figure 4D**).

We continued to explore whether other subject characteristics were associated with cellular *UCP1* expression (**Table S4B**). Strikingly, the only phenotypical feature which was significantly associated with any aspect of cellular UCP1 levels was AGFR, which was negatively correlated to both basal and stimulated deep neck UCP1 (P = 0.012, R^2^ = 0.20, and P = 0.006, R^2^ = 0.23 respectively) (**Figure S4B**, and data not shown). This emphasize a possible connection between overall fat distribution and thermogenic potential of progenitor as well as mature adipocytes. It should however be noted that there was no difference between UCP1 levels or inducibility between the upper and lower quartile, if dividing the dataset based on WHR (**Figure S4C**). Thus, even the individuals with the most pronounced abdominal fat accumulation possessed brown fat precursors in the deep neck fat depot. Taken together these results indicate that despite an observed inverse effect of age, abdominal obesity, and impaired glucose tolerance on UCP1 expression in deep neck fat of adult humans, only the ratio between upper and lower fat deposition seem to negatively affect the abundance or determination of the brown fat precursor cells residing within the tissue, abide not to the extent where such precursors were absent. Thus, brown fat precursor cells appear to remain present during obesity and metabolic derangement, suggesting that reduced thermogenic function as previously observed by others and as shown at a molecular signature level in the current study, is related to a lack of thermogenic programming of the progenitor cells in vivo.

## Discussion

We demonstrate that with increasing BMI, deep neck BAT of adult humans possesses a reduced expression of a subset of thermogenic genes, which normally distinguish it from WAT. Importantly, we find that most markers of BAT identity remained unchanged including a novel distinct immunological signature of BAT, which is retained in obesity along with brown adipose progenitor cells which demonstrated thermogenic capacity when differentiated in vitro. The loss of functionally competent BAT was associated with impaired metabolic health as represented by correlations between decreased *UCP1* mRNA levels and increased abdominal fat distribution as well as impaired glucose metabolism and a dramatic loss of the thermogenic phenotype in BAT of patients with T2DM. Whitening of BAT has been described previously in obesity models of mice where a reduction of VEGF-A-mediated vascularization was identified as a regulatory factor^26^, and in a genetic rat model of type 2 diabetes and obesity^27^. The BAT whitening in mice was characterized by increased accumulation of lipid droplets^26^ as well as reduced mitochondrial concentrations and impaired glucose uptake^27^, suggesting remodeling of BAT into a more WAT-like phenotype. Additional studies in mice, using thermoneutrality to reduce adrenergic activation of BAT, have reported similar results in terms of altered BAT morphology, with an accumulation of larger lipid droplets and a reduced BAT activity, although whitening has not been the target of investigation in these studies^24,25^. Our data suggest that BAT identity is retained in terms of presence of thermogenic progenitors, a subset of BAT unchanged markers as well as the newly defined BAT immune signature. This indicate that although a thermogenic signature is downregulated, BAT remains BAT, but with some adaptations. These adaptations also include changes in cytoskeleton organization, possibly reflecting increased lipid accumulation as this has been shown to be associated^84^. Whether the loss of a thermogenic signature and the gain of a cytoskeleton signature are coupled, or independent events is currently elusive. Waist/hip ratio (WHR) is an established surrogate for fat distribution^86^ and can thus be utilized to estimate abdominal obesity. We here found that increased WHR is negatively correlated with decreased *UCP1* mRNA expression independently of both age and sex and our multivariate analysis indicated that WHR was the driving factor of the observed associations between *UCP1* and glucose homeostasis in our dataset. These data support previous indications that the function of BAT is clearly integrated in the metabolic heath of adult humans^1^. It furthermore raises the idea of a coordinated regulation of BAT and WAT across depots, possibly mediated by circulating factors. The last decade has underscored the dynamics of WAT which can acquire BAT-like properties in response to increased sympathetic activity^20,87^ or by directly enhancing Pparγ signalling^21^. In fact, it has been proposed that adult human BAT should be classified as beige (also known as brite) fat^18,19,88^. Our data leverage the understanding of this matter by raising the possibility that the beige-like morphology of adult human BAT is the result of an age-dependent dormancy, further accelerated into reduced thermogenic capacity by obesity. The mechanism for this is currently unknown but might be related to a reduced vascularization as has been shown to regulate whitening, browning and beiging in mice^26,89^. From a larger perspective, our study illustrate that the plasticity of adipose tissue is not limited to WAT accumulating BAT-like heat-producing properties in response to cold but could also be exemplified by BAT gaining WAT-like energy-storing properties in association with obesity. Our data further suggests that other distinct functions in addition to thermogenic capacity defines BAT, including diverse immunological properties. In conclusion, we propose a novel perspective on the predefined fate of different adipose types and suggest a whole-body interactive regulation of adipose tissue function in adult humans during obesity.

## Methods

### Study design and participants

35 patients scheduled for surgery due to benign goiter were included in the main study, and 37 were additionally included in the biopsy part only, between July 2014 and October 2016 through the outpatient clinics of the Otorhinolaryngology, Head and Neck Surgery and Audiology Departments of Rigshospitalet / Gentofte Hospital and the Department of Otorhinolaryngology and Maxillofacial Surgery, Zealand University Hospital Køge, Denmark. Apart from thyroid malignancy and inability to provide informed consent there were no specific exclusion criteria if patients were considered eligible for surgery. The 35 participants in the main study took part in an additional test day, which included a range of blood samples (table S1A), a DXA-scan and an OGTT at the Centre for Physical Activity Research (CFAS)/ Centre for Inflammation and Metabolism (CIM), Rigshospitalet. Two of the participants of the main study were excluded after the supplemental testing, in one case because surgery was cancelled, and in the other because of acute rescheduling, which made collection of biopsies unachievable. All subjects provided written informed consent prior to participation. The Scientific-Ethics Committees of the Capital Region of Denmark approved the study protocol and amendments, journal number; H-1-2014-015 and the study was performed in accordance with the Helsinki declaration. Basic health information was obtained from all subjects and included a questionnaire about concomitant medical disorders, use of medications or dietary supplements, physical activity level, menopausal status for the women and recent weight changes. Basic health measurements included blood pressure (BP), heartrate (HR), and anthropometrics in the form of body weight (BW) and height (H), and waist and hip circumference measured two cm above the umbilicus with the participant in a relaxed, exhaled, upright position and with legs together at the widest point respectively. Waist-hip measurements were unintendedly not obtained in 14 participants who participated in the biopsy part only. Body fat percentage and android and gynoid fat percentage was assessed using a Dual Energy X-ray Absorptiometry (DXA) scanner (Lunar Prodigy Advance, GE healthcare, Madison; WI, USA). The supplemental test day was attempted scheduled as close to the day of surgery as possible, but for logistical reasons including rescheduling and cancelations of surgery, the number of days between supplemental testing and surgery was unfortunately substantial in a few cases (median number of days = 16, range 1-285 days). In all cases non-invasive measurements (weight, height and W/H, BP and HR) and any changes in medical conditions, medications or lifestyle was reported at the time of surgery.

### Blood sampling and Oral Glucose Tolerance Test (OGTT)

OGTTs were performed in the morning after an overnight (12-hour) fast. Participants were instructed to refrain from alcohol and caffeine intake for 24 hours and from vigorous physical activity 48 hours prior to commencement of the test. Participants were instructed to take habitual medications in the morning except for antidiabetics. One subject with type 2 diabetes, however, displayed a very high blood glucose level upon arrival, wherefore insulin treatment at usual dose was initiated before continuation of the test-day after fasting samples were obtained. Fasting blood samples (table S1) were followed by ingestion of 83 grams of dextrose monohydrate dissolved in 293 mL water. Blood for glucose measurements was collected in Vacuette sodium fluoride containing glucose tubes and blood for C-peptide and insulin measurements in lithium heparin tubes at time points: 10, 20, 30, 60, 90 and 120 minutes after glucose ingestion. Samples were immediately placed on ice and spun before being analyzed at the Department of Clinical Biochemistry, Rigshospitalet. OGTT participants were classified as having NGT, IGT, or type 2 diabetes according to the definition from the World Health Organization (WHO), on the basis of blood glucose levels while fasting and at the two-hour timepoint^90^. The homeostatic model assessment of insulin resistance (HOMA-IR1) index for insulin resistance was calculated from fasting glucose and insulin using the formula (FPI x FPG)/22.5 ^91^.

### Subcutaneous and supraclavicular fat biopsies

Surgery was conducted according to standard procedures by experienced surgeons at one of the three involved surgical departments. The abdominal subcutaneous biopsies were obtained using a modified version of the Bergström needle biopsy procedure as previously described ^92^ after induction of general anesthesia and immediately prior to initiation of surgery. Briefly an incision of 0,5-1 cm was made a few centimeters below and lateral to the umbilicus and the needle was inserted with application of vacuum. The supraclavicular deep neck biopsy was collected by the surgeon from the deep neck fatty tissue either at the lymph node level 4, between the 2 heads of the musculus sternocleidomastoideus or at lymph node level 6 frontally of the thyroid gland depending on the specific type of surgery. The biopsy was obtained from the preexisting surgical incision using scalpel and scissor without application of additional surgery. Biopsies were divided into 2 pieces for the following treatment; one piece was snap frozen in liquid nitrogen and directly moved to a −80-degree freezer at Køge Hospital or transported to the CFAS/ CIM laboratory on dry ice and then stored in a freezer at −80°C. The other part of the biopsy was placed in cell culture media (DMEM/F12 with 1% streptomycin / penicillin) and transported on ice to the CFAS/CIM laboratory for immediate isolation of preadipocytes. On two occasions, the subcutaneous biopsy was collected in local anesthesia (2 ml lidocaine 20 mg/ml) at the CFAS/ CIM laboratory, respectively 14 and 10 days prior to the operation and thus collection of the paired supraclavicular sample. This was due to postponement of the planned operation affecting the possibilities of the study investigators to be present. On one occasion no tissue samples were obtained due to lack of liquid nitrogen, one sample-set was excluded due to mix-up of tubes, and in 16 cases a sufficiently sized subcutaneous biopsy for both tissue analyses and cell culture isolation could not be obtained, in which case cell cultures were prioritized.

### Isolation, culture, and differentiation of human adipogenic progenitor cells

Adipogenic progenitor cells were isolated from the stromovascular fraction of the biopsies on the day they were obtained. Biopsies were collected in DMEM/F12 (Gibco) with 1% penicillin / streptomycin (PS; life technologies) and tubes were kept on ice during transport from the operating room to the cell-lab. Biopsies were digested in a buffer containing 10 mg collagenase II (C6885-1G, Sigma) and 100 mg BSA (A8806-5G, Sigma) in 10 ml DMEM/F12 for 20-30 minutes at 37° C while gently shaken. Following digestion, the suspension was filtered through a cell strainer (70 μm size) and cells were left to settle for 5 minutes before the layer below the floating, mature adipocytes was filtered through a thin filter (30 micron). The cell suspension was centrifuged for 7 min at 800 g and the cell pellet was washed with DMEM/F12 and then centrifuged again before being resuspended in DMEM/F12, 1% PS, 10% fetal bovine serum (FBS) (Life technologies) and seeded in a 25 cm^2^ culture flask. Media was changed the day following isolation and then every second day until cells were 80% confluent, at this point cultures were split into a 10-cm dish (passage 0). Cells were expanded by splitting 1:3. Cells from Passage 1-3 were seeded for gene expression experiments in proliferation media consisting of DMEM/F12, 10 % FBS, 1 % PS and 1 nM Fibroblast growth factor‐acidic (FGF‐1) (ImmunoTools). Cells were grown at 37° C in an atmosphere of 5% CO2 and the medium was changed every second day. Adipocyte differentiation was induced two days after preadipocyte cultures were 100% confluent by addition of a differentiation cocktail consisting of DMEM/F12 containing 1% PS, 0.1 μM dexamethasone (Sigma‐Aldrich), 100 nM insulin (Actrapid, Novo Nordisk or Humulin, Eli Lilly), 200 nM rosiglitazone (Sigma‐Aldrich), 540 μM isobutylmethylxanthine (Sigma‐Aldrich), 2 nM T3 (Sigma‐Aldrich) and 10 μg/ml transferrin (Sigma‐Aldrich). After three days of differentiation, isobutylmethylxanthine was removed from the cell culture media, and after an additional three days rosiglitazone was removed from the media for the remaining 6 days of differentiation. On the 12^th^ day of differentiation the media was changed to DMEM/F12, 1% PS for 2 hours before stimulation with 1 μM norepinephrine (NORadrenalin SAD solution 1 mg/ml) or a control solution of isotonic saline (sodium chloride SAD 9 mg/ML) with the addition of the preservative of the NORadrenalin SAD solution, sodium metabisulfit (Na_2_SO_2_O_5_) (Merck). Cells were stimulated with NE / control treatments for 4 hours before being harvested using TRizol (Invitrogen) and were stored on −80° C until analyses were performed. The degree of cell differentiation was evaluated based on a combination of subjective evaluation of the amount of accumulated lipid droplets (% of culture) on the day of stimulation and on FABP4 mRNA expression (Figure S3A, B and C). For one of the cultures no visible estimation had been registered and thus only FABP4 expression was used for the estimation in this cell ID. 26 (12 supraclavicular and 14 subcutaneous) cell IDs were excluded following harvest due to detection of mycoplasma infection (m. Hyorhinis) confirmed by qPCR.

### RNA isolation and qPCR

Total RNA isolation from adipose tissue biopsies was performed using TRizol reagent according to the manufacture’s protocol. RNA was dissolved in nuclease‐free water and quantified using a Nanodrop ND 1000 (Saveen Biotech). Total RNA (0.25μg) was reverse‐transcribed using the High Capacity cDNA Reverse Transcription Kit (Applied Biosystems). cDNA samples were loaded in triplicate and qPCR was performed using Real Time quantitative PCR, using the ViiA^tm^ 7 platform (Applied Biosystems). Relative quantification was conducted by either SYBRgreen fluorescent dye (Applied Biosystems) or TaqMan Gene Expression Assays (Applied Biosystems). All procedures were performed according to the manufacturer’s protocol. Target gene mRNA expression was normalised to the reference gene PPIA and calculated based on the delta-delta method relative to subcutaneous fat. Primer sequences were UCP1: Hs00222453_m1 (pre-designed TaqMan assay, Life Technologies), PPIA: Forward: acgccaccgccgaggaaaac, Reverse: tgcaaacagctcaaaggagacgc.

### RNA sequencing

RNA was isolated as described above. Ribo-Zero® rRNA Removal Kit was utilized for removal of ribosomal RNA. RNA sequencing was performed by the company BGI. Sequencing reads were mapped to the human reference genome (version hg19) using STAR ^93^. Tag directories were generated using HOMER^94^ and exon reads were counted using iRNA-seq ^95^. All downstream analysis was performed using the software R (ref to r). Genes that did not exceed more than 10 reads in any of the samples were removed from the dataset. Normalization and identification of differentially expressed genes was performed using DESeq2 ^96^. Counts were normalized for sequencing depth/library size and for the length of target transcript (reads pr. kilobase, RPK). PCA and clustering were performed on regularized log transformed counts. For all analysis, DE genes are defined as FDR < 0.05. No lower limit for fold change between the examined conditions were specified for DE genes. To predict the contribution of thermogenic phenotype in each sample, the computational tool ProFAT (REF) was employed on raw reads. One sample was excluded as it differed greatly from all other AT samples in terms of gene expression.

Differentially expressed genes were inserted into the web-based online tool David Annotation Tool (https://david.ncifcrf.gov/tools.jsp) to obtain gene ontologies.

### Statistical analyses

Statistical analyses were performed with SAS 9.4 (SAS Institute Inc. 2013. *SAS® 9.4 Statements: Reference*. Cary, NC: SAS Institute Inc.) for clinical correlations with qPCR analyses, whereas RNA-seq statistical analyses were performed using the R package DESeq2^96^. Unless stated otherwise data is presented as means and SEM, with or without visualization of individual data-points. Associations between UCP1 mRNA expression and continuous variables were explored using uni- and multivariate OLS regression with manual backwards elimination of independent variables for the multivariate models. UCP1 values were transformed using log10 to attain normal distribution when assessing associations. Differences in gene expression between groups were assessed by Mann Whitney U test for unpaired comparisons and Wilcoxon matched pairs signed rank test for paired comparisons. The Kruskal–Wallis one-way ANOVA or regular one-way ANOVA was applied for comparisons between 3 or more groups for tissue samples, depending on whether samples displayed a normal distribution, while a repeated measure mixed models ANOVA with Bonferroni correction was applied for cellular group comparisons. A p-value below 0.05 was considered statistically significant.

## Author Contributions

CS supervised the study. CS, NZJ, SN, BKP, SM, PH, CHH: hypothesis generation, conceptual design, data analysis, and manuscript preparation. NZJ, TWS, SN, MWA, VHJ, MS, LP, IF, RS, ESA conducting experiments and data analysis. All authors edited and approved the final manuscript.

## Acknowledgements

The authors thank the study participants and the staff at the clinical departments. We also thank Pierre Nourdine Bouchelouche from the Department of Clinical Biochemistry, Zealand University Hospital of Copenhagen assistance for the use of their facilities. The Centre for Physical Activity Research (CFAS) is supported by a grant from TrygFonden. During the study period the Centre for Inflammation and Metabolism (CIM) was supported by a grant from the Danish National Research Foundation (DNRF55). The Novo Nordisk Foundation Center for Basic Metabolic Research (http://www.metabol.ku.dk) is supported by an unconditional grant from the Novo Nordisk Foundation to University of Copenhagen. The study was further supported by a research grants from the Lundbeck foundation (CS), the Danish Diabetes Academy supported by the Novo Nordisk Foundation (NZJ) and Novo Nordisk A/S (CS).

## Disclosure of limited availability of biological material and RNA sequencing data

We hereby disclose that the availability of biological material including the human cell cultures and RNA sequencing data is dependent on specific permission from the Danish Data Protection Agency and on the researcher’s adherence to the specifications of this permission.

## Notes

### Competing Interest Statement

The authors have declared no competing interest.

