## Supplemental tables and figures for "Thermogenic genes are blunted whereas brown adipose tissue identity is preserved in human obesity"

### Supplemental information

**Table S1A. Plasma variables examined in relation to UCP1 expression.**

| | Women N = 25 (26) | Men N = 7 | Total N = 32 (33) | Reference interval | Significant correlation with UCP1 |
|---|---|---|---|---|---|
| **HbA1c (mmol/mol)** | 34 (26 – 43) | 37 (33 – 52) | 34 (26 - 52) | < 48 | Y (P < 0.05) |
| **FPG (mmol /l)** | 5.0 (3.9 – 6.2) | 5.1 (4.2 – 10.1) | 5.0 (3.9 – 10.1) | 4.2 – 6.3 | N |
| **PGT120 (mmol/l)** | 5.9 (4.9 – 11.0) | 6.7 (4.1 – 13.5) | 6.3 (4.1 – 13.5) | < 7.8 | Y (P < 0.05) |
| **TSH (x$10^{-3}$ IU / l)** | 0.7 (0.0 – 9.0) | 0.9 (0.2 – 3.0) | 0.86 (0.0 – 9.0) | 0.65 – 4.80 | N |
| **Thyroxin (T4) (pmol/l)** | 15.7 (9.7 – 22.1) | 14.1 (11.3 – 17.6) | 15.5 (9.7 – 22.1) | 14.0 – 23.0* | N |
| **Triiodothyronine (T3) (pmol/l)** | 5.0 (3.9 – 6.0) | 4.9 (2.1 -5.5) | 4.9 (2.1 – 6.0) | 4.1 – 6.9* | N |
| **TC (mmol/l)** | 4.6 (3.6 – 7.3) | 5.1 (2.9 – 5.8) | 4.6 (2.9 – 7.3) | < 5.0 | N |
| **LDL (mmol/l)** | 2.7 (1.8 -5.4) | 3.6 (1.1 – 4.3) | 2.8 (1.1 -5.4) | < 3.0 | N |
| **HDL mmol/l)** | 1.5 (0.8 – 2.5) | 1.1 (1.0 – 1.5) | 1.49 (0.8 – 2.5) | >1.0 | N |
| **TG (mmol/l)** | 0.9 (0.5 – 2.5) | 1.0 (0.7 – 3.2) | 0.96 (0.5 – 3.2) | < 2.0 | N |
| **TLC ($10^9$/l)** | 7.3 (3.3 – 9.8) | 5.6 (4.0 –11.0) | 6.05 (3.3 – 11.0) | 3.5 – 8.8 | N |
| **TNC ($10^9$/l)** | 3.2 (1.5 – 7.1) | 3.4 (2.3 – 6.2) | 1.9 (1.5 – 7.1) | 1.6 – 5.9 | N |
| **HsCRP (mg/l)** | 2.0 (0.3 – 9.6) | 0.6 (0.2 - 6.8) | 3.3 (0.2 – 9.6) | NA | N |

Data are median and range. (N) = number of participants in cell study. Abbreviations: HbA1c = glycated hemoglobin A1c, FPG = fasting plasma glucose, PGT120 = plasma glucose at 120 min, following an oral glucose load, TSH = thyroid stimulating hormone, TC = total cholesterol, LDL = low-density lipoprotein, HDL = high density lipoprotein, TG = triglyceride, TLC = total leucocyte count, TNC = total neutrophil count, HSCRP = high sensitive C-reactive protein. UCP1 = uncoupling protein1, N = no, Y = yes. NA = not available * Age and sex-dependent

**Table S1B. Variables significantly correlated with UCP1.**

| Variable | β-coefficient | P-value | SE | t value | 95% CI |
|---|---|---|---|---|---|
| **WHR** | -4.99 | 0.011 | 1.9 | -2.64 | -8.80 -1.19 |
| **Age** | -0.03 | 0.016 | 0.016 | -2.50 | -0.07 -0.01 |

Total N = 53. Model: Root mean squared variance (RMSE) = 1.177, F-value = 11.87, P < 0.0001, adjusted $R^2$ = 29.0. Correlations were assessed between log10 [UCP1] and subject characteristics by a multivariate ordinary least square (OLS) regression analyses applying manual backwards elimination of the independent variables based on lowest effect size and significance level in the following order; body mass index (BMI),sex, and daily mean outdoor temperature one week prior to surgery. WHR = waist/ hip ratio.

**Table S1C. Clinical characteristics explaining UCP1 variation.**

| Variable | β-coefficient | P-value | SE | t value | 95% CI | |
|---|---|---|---|---|---|---|
| **AGFR** | -1.16 | 0.017 | 1.84 | -1.38 | -2.9 | 0.57 |
| **Age** | -0.03 | 0.061 | 0.018 | -1.85 | -0.07 | 0.002 |
| **PGT120** | -0.16 | 0.056 | 0.084 | -2.0 | -0.34 | 0.005 |

Total N = 31. Model:Root mean squared variance (RMSE) = 0.968, F-value = 7.21, P = 0.001, adjusted $R^2$ = 38.0, coefficient of variation (CV, or relative standard variation . Correlations were assessed between log10 [UCP1] and subject characteristics by a multivariate ordinary least square (OLS) regression analyses applying manual backwards elimination of the independent variables based on lowest effect size and significance level in the following order; body mass index (BMI), fasting plasma glucose (FPG), hemoglobin A1c (HbA1c), and sex. AGFR = android/ gynoid fat ratio, PGT120 = plasma glucose 120 min into the OGT

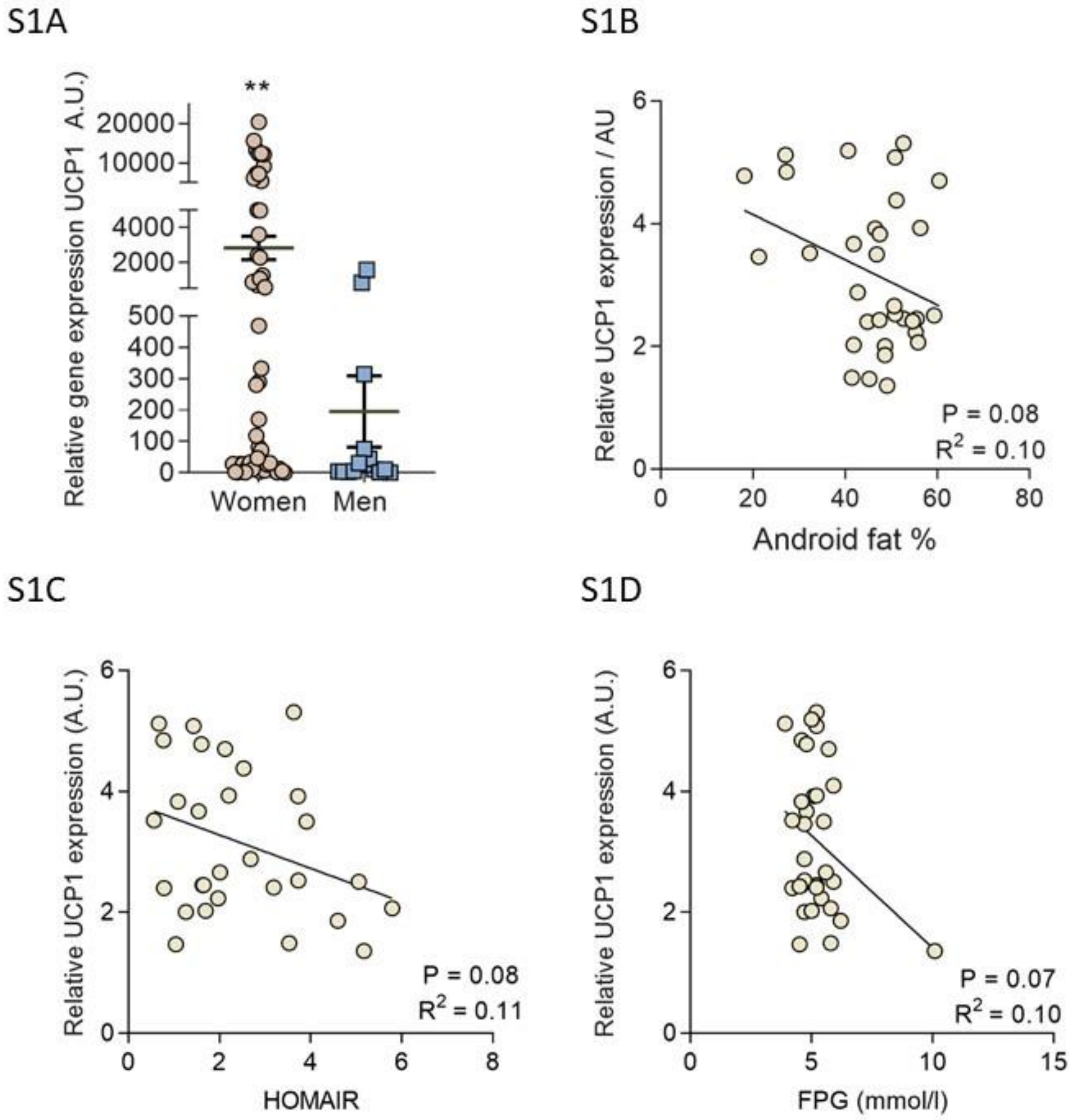


**Figure S1. Clinical characteristics and UCP1 expression**. **A)** Relative UCP1 mRNA expression n deep neck BAT of men (n = 15) and women (n = 53). * = P = 0.008 (two-tailed). Between group differences were calculated using a Mann Whitney U test. Actual mean difference = -214.8 **B)** Correlation between log10 transformed deep neck UCP1 expression and android fat percentage (AF %); β-coefficient = -0.37, 95% CI (-0.078 to 0.004), F = 3.3, and **C)** homeostatic model assessment of insulin resistance (HOMAIR) (N = 29); β-coefficient = - 0.28, 95% CI (-0.059 to 0.033), F = 3.4 **D)** fasting plasma glucose levels (FPG) (N = 33), β-coefficient = - 0.37, 95% CI (-0.78 to 0.039), F = 3.4. Data were analyzed using univariate linear regression analyses and are presented with unadjusted P-values. N= 32, unless specified otherwise. AU = arbitrary units. +2 were added to Log10 transformed UCP1 values for graphical presentation of correlations

**Table S2A Top 10 contributing genes to PC1 (44.22 %) and PC2 (18.6%) in PCA, Figure 2B.**

| PC1 | PC2 |
|---|---|
| UCP1 | PTX3 |
| SELE | FOSB |
| HMGCS2 | CLEC4M |
| APOB | SELE |
| OGDHL | FOS |
| FOSB | SERPINE1 |
| IL6 | TIPARP |
| CXCL8 | MS4A1 |
| SLC36A2 | FOSL1 |

**Table S2B. top 20 BAT selective genes according to log2FC and P-values respectively**

| Symbol | Delta-W-B | log2FC_W.vs._B | Padj_W.vs.B | Symbol | Delta-W-B | log2FC_W.vs._B | Padj_W.vs.B |
|---|---|---|---|---|---|---|---|
| *UCP1* | -6565,27 | -9,46784 | 1,37E-07 | *WNT5A* | -87,2423 | -3,28082 | 4,65176E-17 |
| *OGDHL* | -146,861 | -8,968 | 7,01E-06 | *CA13* | -57,8579 | -2,95908 | 1,9802E-15 |
| *MZB1* | -129,929 | -8,43827 | 4,69E-06 | *CYSLTR2* | -315,584 | -4,82376 | 6,68149E-12 |
| *LHX8* | -22,809 | -8,02185 | 3,69E-06 | *HHIPL2* | -94,9557 | -5,94659 | 1,8814E-11 |
| *ZDHHC19* | -82,9914 | -7,90681 | 0,0002 | *CYP1A2* | -200,564 | -7,61741 | 1,58409E-10 |
| *IL6* | -1585,33 | -7,82095 | 8,76E-07 | *SCEL* | -45,8623 | -3,71296 | 1,58409E-10 |
| *SELE* | -606,35 | -7,65752 | 4,51E-05 | *CPN2* | -99,0004 | -6,60928 | 2,18246E-10 |
| *CYP1A2* | -200,564 | -7,61741 | 1,58E-10 | *SLC27A6* | -53,4118 | -4,77466 | 4,9777E-10 |
| *FOSB* | -1706,9 | -7,56821 | 7,99E-05 | *CNTN4* | -37,4473 | -2,682 | 8,86978E-10 |
| *IGLL5* | -1119,56 | -7,47822 | 4,14E-06 | *FBXO40* | -16,6341 | -3,22754 | 9,63299E-10 |
| *FUT3* | -43,5647 | -7,24091 | 0,0090 | *FBP2* | -95,5716 | -6,30061 | 2,83337E-09 |
| *CXCL1* | -53,7883 | -7,09264 | 3,67E-05 | *LEPR* | -1009,18 | -2,48962 | 1,0003E-08 |
| *NTS* | -173,257 | -7,04683 | 9,48E-07 | *CKMT1B* | -572,168 | -6,94982 | 1,26132E-08 |
| *FCRL5* | -23,8092 | -7,03225 | 0,00045 | *PTX3* | -2010,46 | -6,8506 | 1,28453E-08 |
| *TBATA* | -195,674 | -6,96484 | 5,96E-06 | *CMYA5* | -142,227 | -2,14924 | 3,24916E-08 |
| *CKMT1B* | -572,168 | -6,94982 | 1,26E-08 | *CLSTN3* | -675,033 | -3,41889 | 5,8914E-08 |
| *PTX3* | -2010,46 | -6,8506 | 1,28E-08 | *CYP1B1* | -275,843 | -2,40973 | 6,85429E-08 |
| *CKMT1A* | -237,209 | -6,79325 | 2,9E-07 | *SELPLG* | -179,647 | -2,37827 | 7,48973E-08 |
| *IGJ* | -1306,17 | -6,70245 | 7,08E-07 | *RNF175* | -58,8724 | -2,72477 | 1,25982E-07 |
| *CXCL13* | -78,6898 | -6,66899 | 0,0027 | *UCP1* | -6565,27 | -9,46784 | 1,37355E-07 |

Top 20 DE genes with highest expression in BAT, according to log2foldchange WAT vs. BAT log2FC_W.vs.B) (left panel) and lowest adjusted p-values (Padh_W.vs.B) (right panel).

**Table S2C. top 20 WAT selective genes according to log2FC and P-values respectively.**

| Symbol | Delta-W-B | log2FC_W.vs._B | Padj_W.vs.B | Symbol | Delta-W-B | log2FC_W.vs._B | Padj_W.vs.B |
|---|---|---|---|---|---|---|---|
| *SIM1* | 94,89188 | 7,86456 | 2,60422E-05 | *APOB* | 352,0901 | 7,475002 | 5,48164E-19 |
| *APOB* | 352,0901 | 7,475002 | 5,48164E-19 | *AFF2* | 75,8672 | 3,900455 | 4,4848E-17 |
| *EGFL6* | 263,5983 | 7,310409 | 7,73284E-07 | *HOXB7* | 156,5014 | 3,252377 | 1,24232E-16 |
| *PAX3* | 29,8824 | 6,896275 | 0,001488097 | *GALNT13* | 43,99199 | 3,589217 | 4,67427E-14 |
| *MYEOV* | 75,75396 | 6,552332 | 2,06584E-08 | *ANKRD33B* | 151,327 | 2,319326 | 5,38083E-13 |
| *IRX1* | 388,3313 | 5,771029 | 9,5568E-11 | *NPY5R* | 165,8694 | 3,110662 | 1,50708E-12 |
| *SP9* | 13,2373 | 5,450459 | 0,000265672 | *NPY1R* | 916,3012 | 3,163344 | 1,62381E-11 |
| *HOXA13* | 3,641932 | 5,207508 | 9,21737E-05 | *HOXB8* | 30,2898 | 4,3468 | 1,75052E-11 |
| *URAD* | 19,79347 | 4,98962 | 0,0364594 | *PCLO* | 97,6089 | 4,817562 | 2,16159E-11 |
| *GDA* | 13,5879 | 4,853106 | 2,68218E-06 | *HOXC8* | 59,77028 | 3,631669 | 2,81109E-11 |
| *PCLO* | 97,6089 | 4,817562 | 2,16159E-11 | *HOXB6* | 100,2069 | 3,391413 | 3,87941E-11 |
| *NRCAM* | 148,0733 | 4,738043 | 7,39019E-07 | *ERBB4* | 61,31203 | 3,700758 | 6,40018E-11 |
| *HOXA11* | 14,55753 | 4,649184 | 7,0831E-07 | *IRX1* | 388,3313 | 5,771029 | 9,5568E-11 |
| *CSN1S1* | 37,36855 | 4,519589 | 0,004310947 | *DDI2* | 285,1157 | 0,988112 | 4,44572E-10 |
| *HOXB8* | 30,2898 | 4,3468 | 1,75052E-11 | *COL8A1* | 442,131 | 3,205113 | 1,24707E-09 |
| MMP13 | 8,72214 | 4,302133 | 0,001175823 | LPGAT1 | 451,3033 | 1,353974 | 2,79164E-09 |
| KRT16 | 72,09266 | 3,972608 | 0,005086296 | OSGIN2 | 452,6929 | 1,764274 | 2,79164E-09 |
| VGLL1 | 4,079245 | 3,965976 | 0,01393491 | RNF144A | 289,5799 | 1,418872 | 2,90561E-09 |
| AFF2 | 75,8672 | 3,900455 | 4,4848E-17 | SCARB1 | 399,5086 | 1,539562 | 4,79269E-09 |
| HOXC10 | 47,30777 | 3,748158 | 6,08689E-06 | ADK | 138,6282 | 1,121724 | 5,55331E-09 |

Top 20 DE genes with highest expression in WAT, according to highest log2foldchange WAT vs. BAT (log2FC_W.vs.B) (left panel) and lowest adjusted p-values (Padj_W.vs.B) (right panel).

**Table S3. Subject characteristics of participants included in the RNA-seq analyses.**

| Category | Normal weight WAT | Normal weight BAT | Overweight BAT | Obese BAT |
|---|---|---|---|---|
| **Sex (M: W)** | 1:5 | 2:7 | 3:8 | 3:9 |
| **Age (years)** | 52 (23 – 67) | 47 (23 – 67) | 53 (29 -79) | 56 (42 – 70) |
| **BMI (kg/m$^2$)** | 22.5 (17.6 – 24.9) | 22.6 (17.6 – 24.9) | 27.4 (25.1 – 29.8) | 35.1 (30.8 – 42.9) |
| **WC (Cm)** | 78.5 (61 -96) | 75 (61 – 96) | 92.5 (84 – 107) | 110.5 (102 – 133) |
| **WHR** | 0.8 (0.7 -1.0) | 0.8 (0.7 – 1.0) | 0.9 (0.7 – 1.0) | 0.9 (0.8 – 1.1) |
| **T2DM** | N = 1 | N = 1 | N = 1 | N = 2 |

Values are median and range. M = men, W = women. BMI = Body Mass Index, WC = waist circumference, WHR = waist / hip ratio, T2DM = type 2 diabetes. Patients with type 2 diabetes were only included in the profit analyses.

**Table S4. Cell donors for Seahorse experiments**

| | **Normal weight** | **Obese** |
|---|---|---|
| **Sex** | 5F, 0M | 5F, 0M |
| **Age** | 50 +/- 7.5 | 56 +/- 8.3 |
| **BMI** | 23.2 +/- 2.2 | 38 +/-1.0 **** |
| **%lipid droplets in cells** | 66 +/- 18 | 61 +/- 19 |

Data are mean +/- SD. ****P<0.0001, unpaired t-test.

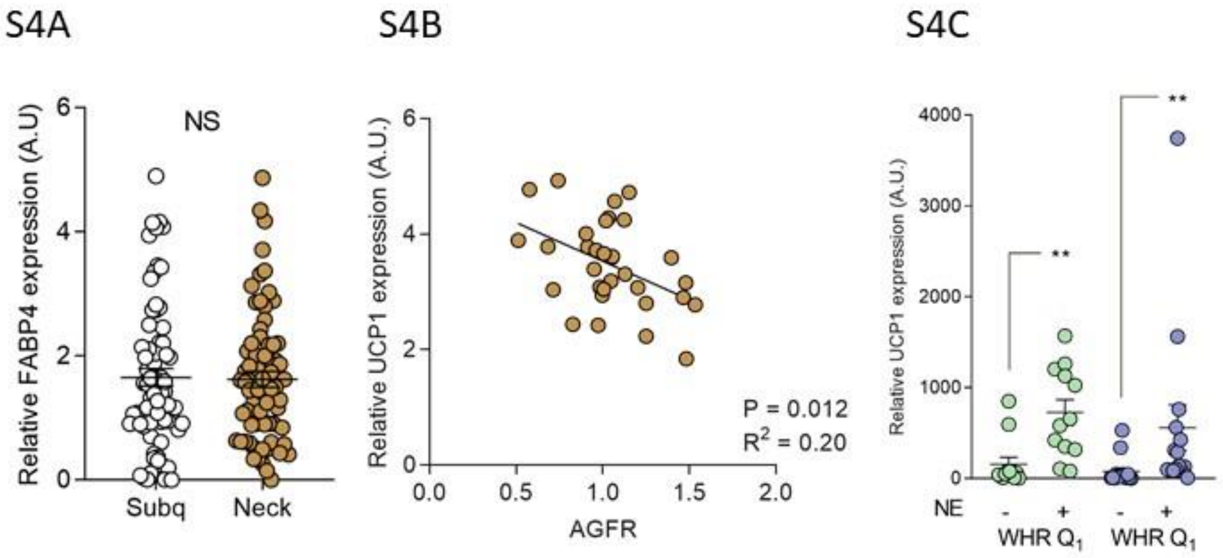


**Figure S4. Isolated in vitro differentiated cells. A)** FABP4 mRNA expression (mean and SEM) in subcutaneous WAT and deep neck BAT unstimulated cells (n = 67 paired IDs). A paired two-sided students t-test was performed. P = 0.94, r = 0.49, t =0.08 df = 68, Mean of differences = - 0.01, 95% CI (-0.28 to 0.26), $R^2$ = 9.505e-005. **B)** Correlation between log10 transformed deep neck adipocyte basal UCP1 expression and android/ gynoid fat ratio (AGFR)), P < 0.012, $R^2$ = 0.20, β-coefficient = - 1.34, 95% CI (-2.34 to -0.33), F = 7.41 **C)** Basal and norepinephrine (NE) induced UCP expression in brown adipocytes derived from patients within the lower ($Q_1$) and upper ($Q_2$) quartile of waist-hip-ratio values. Data was analyzed with a repeated measure mixed models ANOVA with Bonferroni correction. ** = P < 0.01. AU = arbitrary units, NS = not, ctrl = control conditions, NE = norepinephrine, AGRF = android-gynoid-fat-ratio, WHR = waist-hip-ratio.
